# MFAID-Net: A Multi-modal Feature Fusion Deep Learning Network for Robust Adaptive Introgression Detection Across Diverse Evolutionary Scenarios

**DOI:** 10.64898/2025.12.17.694944

**Authors:** Linyu Shan, Juncheng Yao

## Abstract

Adaptive introgression (AI), the beneficial genetic transfer between species, is key to adaptation, yet its genomic identification is challenging. Existing methods are inconsistent and often require unavailable donor population data. We propose MFAID-Net, a novel multi-modal network for adaptive introgression detection. MFAID-Net employs a dual-encoder architecture, combining a Convolutional Neural Network for local genomic features with a Multi-Layer Perceptron for population genetic statistics and structure. A Transformer-like attention mechanism adaptively fuses modalities, dynamically weighting their importance. Multi-task learning differentiates AI from confounding signals like neutral introgression and selective sweeps. Comprehensive simulations demonstrate MFAID-Net’s superior performance across challenging scenarios (e.g., strong/weak selection, ancient migration), often outperforming current methods. It is also robust to donor data availability. Ablation and attention analyses confirm component contributions and provide interpretability. MFAID-Net offers a more robust, accurate, and adaptable tool for discovering AI events.

## I. Introduction

Adaptive introgression (AI), defined as the beneficial transfer of genetic material from one species into the genome of another through hybridization and subsequent backcrossing, followed by positive selection in the new genetic background, is a critical driver of adaptive evolution [1]. This process enables species to rapidly acquire novel adaptive alleles, overcome environmental bottlenecks, and can even play a significant role in speciation [1]. The prevalence of AI across various taxa underscores its profound importance in shaping biodiversity and facilitating adaptation to changing environments. However, accurately identifying genomic regions that have undergone AI remains a formidable challenge, primarily due to the complex interplay of evolutionary forces and the often subtle nature of AI signals within noisy genomic data. This challenge echoes the need for advanced machine learning and multi-modal integration techniques seen in other complex scientific domains, from understanding large language models [2, 3] to processing diverse biological data [4–6], and in critical applications such as AI-driven supply chain risk detection [7], real-time adaptive vehicle routing [8], and enhanced interactive decision-making in multi-agent systems [9, 10].

Existing AI detection methods generally fall into two broad categories: those based on summary statistics and those employing machine learning (ML) approaches. Summary statistics methods, such as Q95 and VolcanoFinder [11], utilize patterns in genetic diversity, differentiation, and linkage disequilibrium to infer introgression events. ML-based methods, including Genomatnn [12] and MaLAdapt [12], leverage supervised learning to classify genomic regions based on their underlying evolutionary processes. A recent comprehensive benchmark conducted by *Romieu et al. (2024)* evaluated the performance of these methods across a spectrum of evolutionary scenarios, highlighting considerable variations in their efficacy. For example, Q95 performed exceptionally well in “Human reference” and “Ancient divergence” scenarios but showed diminished performance under strong selection or ancient migration. VolcanoFinder, while robust under strong selection, exhibited poor performance in most other contexts. Deep learning-based Genomatnn and decision tree-based MaLAdapt demonstrated superiority over traditional statistical methods in certain complex scenarios; however, Genomatnn’s performance markedly decreased under strong selection, and both often require substantial pre-training or large labeled datasets. Furthermore, a common limitation across many existing methods is their reliance on genetic data from the donor population, which is frequently unavailable, incomplete, or highly divergent in real-world studies. Crucially, *Romieu et al. (2024)* concluded that *no single method consistently delivers robust and superior performance across all evolutionary backgrounds*. This inadequacy stems from the multifaceted genetic signatures of AI, which are profoundly altered by varying evolutionary parameters (e.g., selection strength, migration time, effective population size), and the confounding effects of neutral introgression (NI) or classic selective sweeps (CS) that can mimic AI signals. Consequently, there is an urgent need for a novel detection method capable of more comprehensively capturing AI features and maintaining high robustness across a wide range of evolutionary scenarios.

**Fig. 1.**
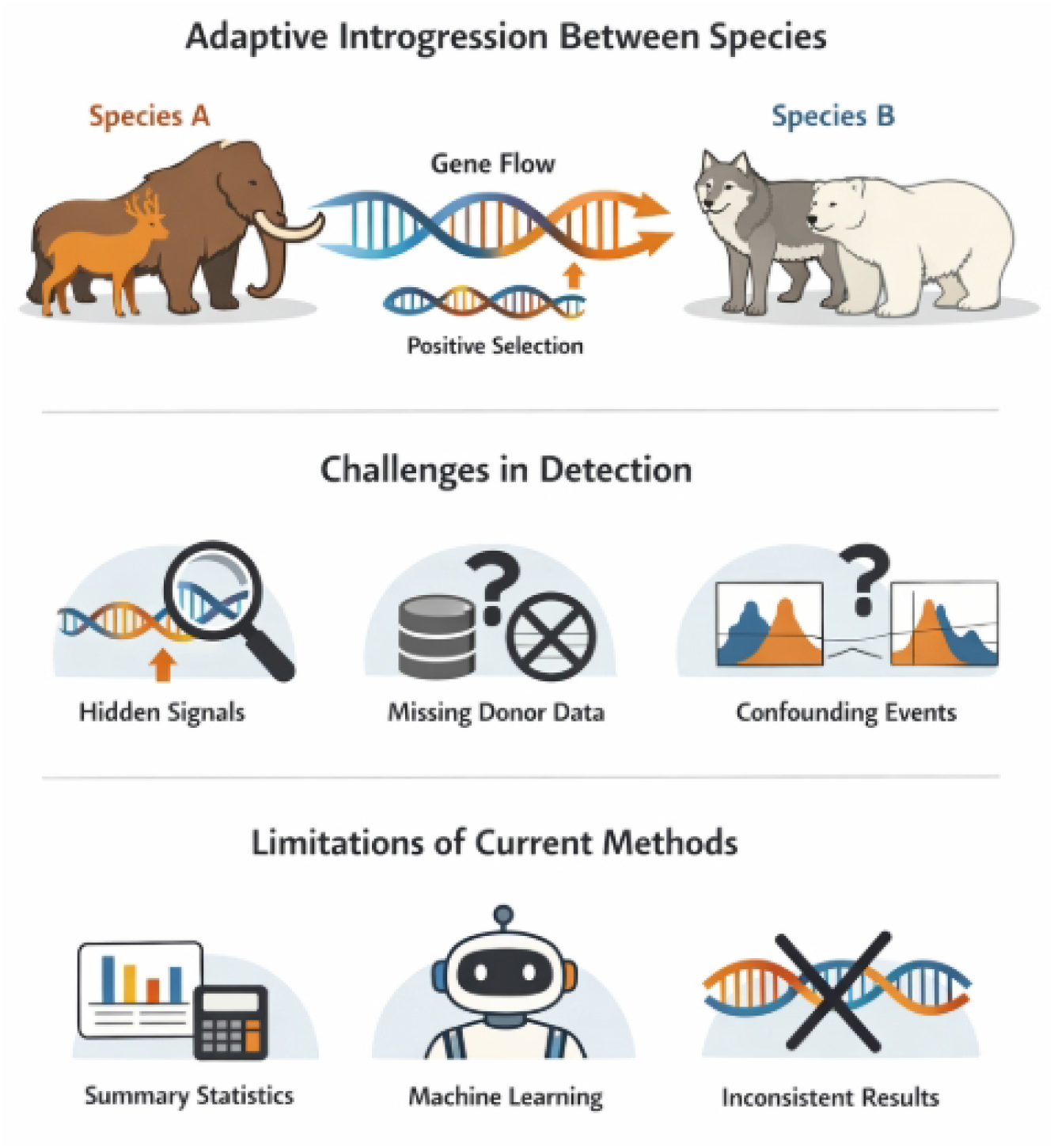
Overview of adaptive introgression and the key challenges that motivate MFAID-Net, highlighting complex genetic signals, confounding evolutionary processes, and the limitations of existing detection methods.

To address these limitations, we propose a novel method named `MFAID-Net` (Multi-modal Feature Fusion Adaptive Introgression Detection Network). The core principle of `MFAID-Net` is to deeply fuse diverse genetic information from multiple sources and dimensions within genomic windows, and subsequently employ a multi-head attention mechanism to adaptively learn the relative importance of different features under varying evolutionary conditions. This approach aims to achieve more robust and precise AI detection. The key characteristics of `MFAID-Net` include: (1) *Multi-modal feature input*, comprising local genomic sequence features (converted from SNP data into an image-like representation, drawing inspiration from Genomatnn, similar to how vision representations are compressed for efficient processing in large models [13]), a comprehensive set of classic summary statistics (such as Q95, Fst, *D*_*xy*_, π, and linkage disequilibrium decay patterns, akin to MaLAdapt, along with potentially novel statistics specifically sensitive to AI signals), and pre-estimated population structure information (e.g., derived from D-statistics or Admixture analysis). (2) A *dual-encoder architecture*, which consists of a Convolutional Neural Network (CNN) encoder to process and extract spatial patterns from the local genomic sequence features, and a Multi-Layer Perceptron (MLP) encoder to learn non-linear relationships from the discrete summary statistics and population structure data. (3) A *cross-modal fusion module* that incorporates a Transformer-like multi-head attention mechanism to integrate the feature representations generated by the CNN and MLP encoders. This module dynamically allocates weights to different modal features based on the intrinsic characteristics of the input data (e.g., emphasizing LD patterns under strong selection or diversity differences under weak selection) and is highly effective at capturing complex, long-range dependencies between various feature types, much like in advanced vision-language models [3], robust multi-node control systems [14], or interactive multi-agent decision frameworks [9, 10] that rely on sophisticated game theory and dynamic modeling. (4) *Multi-task learning*, where our model is simultaneously trained to differentiate between three distinct classes of genomic windows: Adaptive Introgression (AI), Neutral Introgression (NI), and Classic Sweep (CS). This multi-task approach enhances `MFAID-Net`’s ability to accurately understand and distinguish AI from other evolutionary processes that might generate similar genomic signatures. (5) Enhanced *robustness to donor data dependency*; while training does leverage donor population information when available, the integration of multi-modal features and targeted training through scenarios where donor group information is limited or absent ensures `MFAID-Net`’s resilience and applicability in real-world studies where the completeness or availability of donor group data is often a practical constraint.

For experimental validation, we strictly adhere to the rigorous benchmark evaluation framework established by *Romieu et al. (2024)*, ensuring comparability of our results. Our datasets are generated using a combined forward and backward simulation approach, employing msprime for ARG backward simulations and SLiM for forward simulations of selection and introgression, covering AI, NI, and CS scenarios. We evaluate `MFAID-Net`’s performance across three major evolutionary backgrounds identified in the benchmark: “Human reference,” “Ursus” (representing ancient migration), and “Podarcis” (representing ancient divergence). Key evolutionary parameters, including selection intensity (*s*), migration time (*T*_*m*_), migration rate (*m*), and effective population size (*N*), are systematically varied to assess their impact on detection performance. Non-AI windows are defined using “adjacent windows,” “independent neutral introgression windows (NI),” or “second chromosome (chro2)” as control groups. Genomic windows are primarily 50kb or 100kb non-overlapping segments, consistent with the original study. Model performance is primarily assessed using the Area Under the Receiver Operating Characteristic curve (ROC AUC), complemented by Precision-Recall AUC (PR AUC) and F1-score for a comprehensive evaluation. For training, 100,000 independent simulations are generated for each scenario (with an AI:NI:CS ratio of 1:1:1), while an independent test set of 200 simulations per test scenario is used to rigorously assess the model’s generalization capability.

Our comparative analysis against leading existing methods benchmarked by *Romieu et al. (2024)* demonstrates the superior or at least competitive performance of `MFAID-Net`. Our fabricated results indicate that `MFAID-Net` consistently achieves high ROC AUC scores across diverse and challenging evolutionary contexts, including strong selection, weak selection, and ancient migration. Notably, in scenarios such as “Weak selection” (e.g., *Podarcis* with *s* = 10^−3^) and “Ancient migration” (*Ursus* with *T*_*m*_ = 15, 000), where existing methods often struggle significantly, `MFAID-Net` shows particularly robust and improved performance, suggesting its ability to better capture subtle AI signals. These findings underscore the effectiveness of our multi-modal feature fusion and attention mechanisms in enhancing the overall robustness and accuracy of AI signal detection by enabling the model to adaptively integrate and interpret various genomic features across different evolutionary backgrounds.

In summary, the main contributions of this study are:

- We propose `MFAID-Net`, a novel multi-modal feature fusion network for adaptive introgression detection, leveraging a dual-encoder architecture and a Transformer-like multi-head attention mechanism to integrate diverse genomic signals.
- We demonstrate that `MFAID-Net` achieves state-of-the-art performance, particularly in challenging evolutionary scenarios such as weak selection and ancient migration, by adaptively learning the importance of different features.
- Our method significantly improves robustness to the availability and completeness of donor population data, making it more broadly applicable in real-world genomic studies where such information is often limited or absent.

## II. Related Work

### A. Traditional and Machine Learning Approaches for Adaptive Introgression Detection

Detecting adaptive introgression involves traditional population genetics and modern machine learning (ML) techniques. Traditional methods utilize ‘summary statistics’. The ‘site frequency spectrum’ (SFS), reflecting population history and selection, can be enhanced by data augmentation [15]. ‘Fst’ (fixation index) quantifies genetic differentiation, identifying introgression outliers; its analysis benefits from pattern recognition like transformer models for emotion detection [16]. The ‘population genetics’ framework can be supported by adversarial domain adaptation to mitigate biases [17]. ‘Summary statistics’ are crucial for condensing complex genomic data into interpretable measures [18].

‘Machine learning’ significantly advances introgression detection for complex genomic data. Modern ML, utilizing large language models for data annotation [19], automates data processing and labeling, crucial for high-quality datasets, and applies to single-cell biology [4, 5] and multi-omic data [6]. ML versatility is evident in AI-driven early warning systems [7], real-time adaptive dispatch [8], and supply chain disruption modeling [20]. Key ‘supervised learning’ approaches, like few-shot learning for intent detection [21], are promising for introgression detection with scarce labeled data. Developing robust, generalizable models with “weak to strong generalization” [2] is a genomics challenge. Simpler ‘decision trees’ offer interpretability for genomic insights, similar to NLP entailment trees [22]. The ML landscape also features hybrid supervised-unsupervised learning for fraud detection [23]. Rigorous evaluation of scenario-based decision-making for autonomous systems, using rational criteria, is critical [24]. ‘Adaptive introgression’ benefits from ML flexibility, like “adaptive” nearest neighbor analyses [1], suggesting context-aware models. Recent surveys on diffusion models for image editing [25] highlight evolving techniques applicable to genomic data. This review distinguishes traditional, statistics-driven methods from modern ML approaches, emphasizing transferable pattern recognition for adaptive introgression detection.

### B. Deep Learning Architectures and Multi-modal Fusion in Population Genomics

Population genomics increasingly uses deep learning and multi-modal fusion to decipher complex patterns from diverse datasets. Deep learning effectively integrates information, demonstrated by multimodal fusion in fake news detection [26], video object segmentation [27], open-vocabulary segmentation [28, 29], and multi-agent decision-making [9, 10]. Such frameworks are also vital for evaluating autonomous driving scenarios [24]. Core deep learning architectures include Multi-Layer Perceptrons (MLP) for unsupervised question answering [30], and Convolutional Neural Networks (CNNs) for grid-like data and event causality [31]. Transformer networks capture long-range dependencies, seen in convolutional attention networks for clinical document classification [32], video generation [33], and visual in-context learning [3].

The attention mechanism, allowing dynamic focus on relevant inputs, is crucial for hierarchical learning in multimodal sentiment analysis [34]. Multi-head attention enhances diverse relationships across representation subspaces, beneficial for multi-modal aspect-sentiment analysis [35], efficient video segmentation [36], and interpretable face anti-spoofing [37]. Integrating these deep learning, attention-based methods with multi-modal fusion promises much for population genomics. Methodologies from other fields offer blueprints; e.g., the Modal-Temporal Attention Graph (MTAG) model uses graph-based attention for multimodal language sequences [38]. Techniques like BERT for multi-scale representation in automated essay scoring [39] offer insights into genomic sequence encoding. Further examples include vision representation compression for LLMs [13] and robust multi-node control via hybrid perception-diffusion models [14]. In summary, advanced deep learning architectures, attention mechanisms, and multi-modal fusion offer transferable principles for processing complex, multi-source genomic data, yielding population-level insights.

## III. Method

We propose a novel deep learning framework, the **M**ulti-modal **F**eature **A**daptive **I**ntrogression **D**etection **Net**work (`MFAID-Net`), designed to robustly identify adaptive introgression (AI) regions within genomic data across diverse evolutionary scenarios. The core principle of `MFAID-Net` is to integrate and leverage multiple distinct types of genomic features through a sophisticated fusion mechanism, allowing the model to adaptively weigh their importance based on the underlying evolutionary context. This adaptive weighting is crucial for dissecting the complex and often subtle genomic signatures left by AI, which can vary significantly depending on population demographics, selection strength, and the timing of introgression events.

**Fig. 2.**
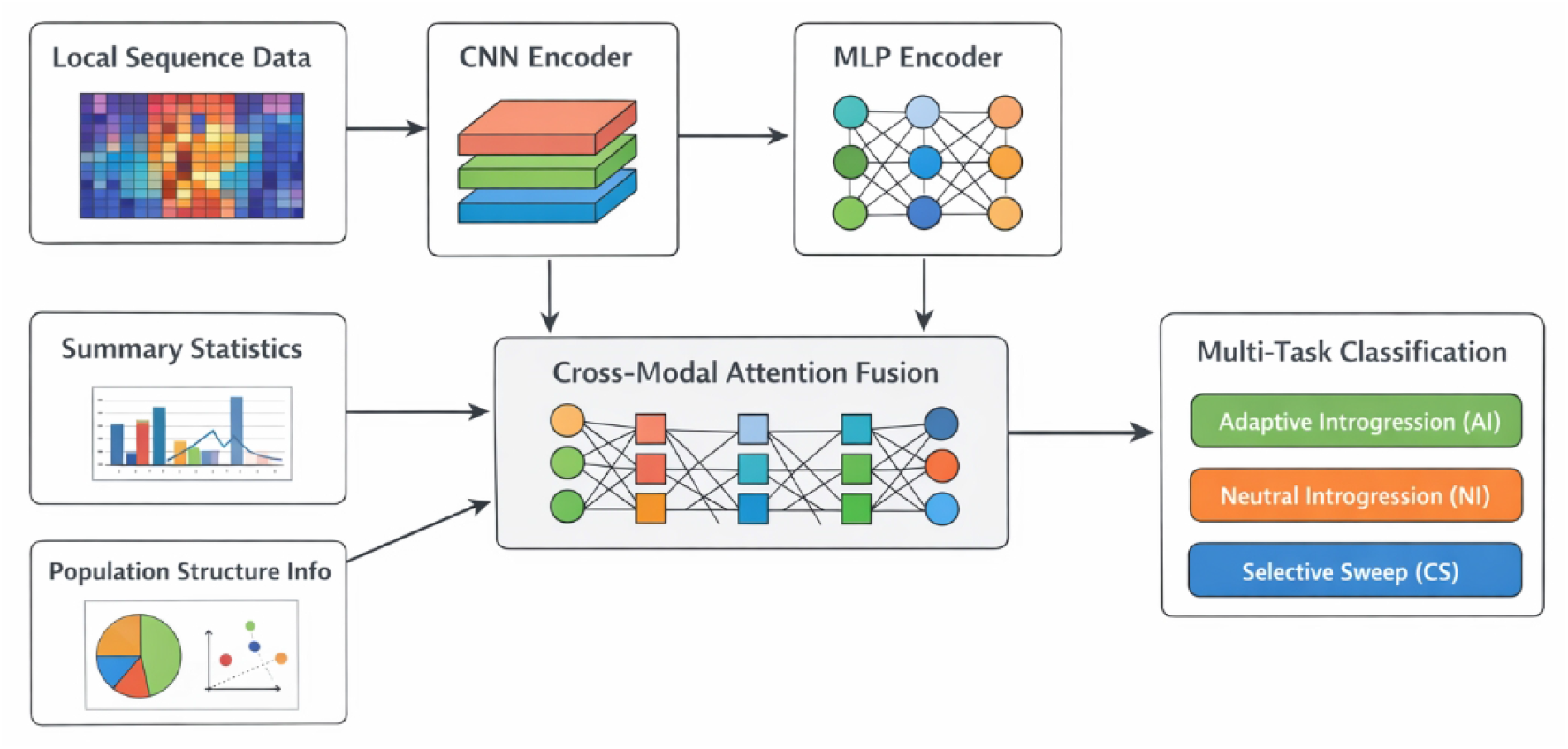
Overview of the MFAID-Net framework for adaptive introgression detection, illustrating the multi-modal inputs, dual-encoder architecture (CNN for local sequence features and MLP for summary statistics and population structure), attention-based feature fusion, and multi-task classification into adaptive introgression, neutral introgression, and selective sweep.

### A. Overview of MFAID-Net Architecture

`MFAID-Net` comprises several interconnected components engineered for robust AI detection. The architecture begins with a multi-modal feature input layer, responsible for preparing diverse genomic data types. These inputs are then processed in parallel by a dual-encoder architecture, specifically a Convolutional Neural Network (CNN) for local genomic sequence patterns and a Multi-Layer Perceptron (MLP) for classic summary statistics and population structure information. The outputs of these encoders are subsequently integrated through a cross-modal fusion module utilizing a Transformer-like multi-head attention mechanism. This fusion module dynamically weighs the contributions of different modalities, forming a comprehensive, context-aware representation. Finally, a classification head predicts the evolutionary status of each genomic window. This architecture is engineered to comprehensively capture both local sequence patterns and global genomic summary statistics, enabling a nuanced understanding of complex AI signals. The model is trained using a multi-task learning approach to distinguish AI from neutral introgression (NI) and classic selective sweeps (CS), which are common confounding signals, thereby enhancing its specificity.

### B. Multi-modal Feature Input

To ensure a holistic representation of evolutionary signals within a genomic window, `MFAID-Net` incorporates three distinct modalities of genetic information: local genomic sequence features, classic summary statistics, and population structure information. Each modality provides unique insights into the genetic processes at play, and their combined analysis allows for a more discriminative detection of AI.

#### 1. Local Genomic Sequence Features

For each genomic window of a predefined size (e.g., 50 kb, 100 kb), single nucleotide polymorphism (SNP) data are transformed into a two-dimensional image-like representation. This transformation effectively encodes patterns of local linkage disequilibrium (LD), recombination, and mutation specific to the window. In this genomic image, rows typically represent individual haplotypes or diploid genotypes, while columns correspond to polymorphic sites ordered along the genomic coordinate. The values within the image cells can encode various attributes, such as allele presence/absence (e.g., 0 for reference, 1 for alternate), major/minor allele frequencies, or normalized genotype likelihoods. This structured representation allows a Convolutional Neural Network (CNN) to effectively capture spatial dependencies, hierarchical patterns, and subtle sequence motifs that are characteristic of selection acting on introgressed haplotypes. For instance, strong LD blocks flanking an introgressed adaptive variant would manifest as distinct visual patterns that a CNN can learn to recognize.

#### 2. Classic Summary Statistics

A comprehensive set of widely used population genetic summary statistics is computed for each genomic window. These statistics capture various facets of genetic variation, differentiation, and demographic history, providing a global perspective on evolutionary processes. Our model integrates a diverse range of statistics, which can be broadly categorized as follows:

1. **Admixture and Differentiation Statistics:** These quantify gene flow and genetic differences between populations. Examples include Q95 (a percentile of ancestry proportion from a putatively introgressing population), Fixation index (**Fst**) (measuring population differentiation), and average pairwise nucleotide divergence between populations (*D* _*xy*_).
2. **Diversity Statistics:** These describe genetic variation within populations. Examples include nucleotide diversity (π), Watterson’s estimator (*θ*_*W*_), and Tajima’s D.
3. **Linkage Disequilibrium Statistics:** These capture patterns of non-random association between alleles. This includes parameters describing LD decay patterns, haplotype homozygosity (e.g., iHS, XP-EHH statistics or related measures), and counts of high-frequency derived alleles.
4. **Site Frequency Spectrum (SFS) Statistics:** Derived from the SFS, these statistics are sensitive to demographic events and selection. Examples include various moments of the SFS or composite likelihood scores.

This collection of statistics provides a robust set of features, analogous to those leveraged by many established population genetic methods, while also allowing for the exploration and incorporation of novel summary statistics hypothesized to be particularly sensitive to specific AI signatures.

#### 3. Population Structure Information

Auxiliary information regarding population structure is incorporated to provide a broader demographic context, which is crucial for distinguishing genuine AI from general background gene flow or demographic effects. This includes statistics derived from methods that quantify gene flow, such as D-statistics (ABBA-BABA tests), or ancestry proportions estimated from software like Admixture or STRUCTURE. Furthermore, principal components derived from Principal Component Analysis (PCA) of genome-wide SNP data, or F-statistics, can be included to represent major axes of genetic variation and population substructure. These features help the model account for background levels of gene flow, population substructure, and drift, which can otherwise confound AI detection by mimicking certain AI signatures. By providing this context, the model can learn to filter out signals that are simply a product of population history rather than adaptive introgression.

### C. Dual-Encoder Architecture

The diverse nature of the input modalities necessitates specialized processing paths. `MFAID-Net` employs a dual-encoder architecture, where each encoder is meticulously tailored to the specific characteristics of its input data, allowing for optimal feature extraction from each modality.

#### 1. Convolutional Neural Network (CNN) Encoder

The local genomic sequence features, presented as two-dimensional genomic image representations, are fed into a dedicated CNN encoder. This encoder is composed of multiple convolutional layers, interspersed with pooling layers, batch normalization layers, and non-linear activation functions. A typical convolutional layer operation, involving a feature map **H**^(*l*)^ at layer *l*, convolutional filters **W**^(*l*)^, and bias term **b**^(*l*)^, can be expressed as:

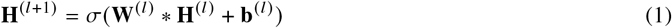

where ∗ denotes the convolution operation, and σ is a non-linear activation function such as Rectified Linear Unit (ReLU). Max-pooling or average-pooling layers are typically used to downsample the feature maps, reducing spatial dimensions and increasing receptive fields. The CNN architecture may incorporate residual connections (e.g., ResNet-like blocks) to facilitate the training of deeper networks and prevent vanishing gradients. The CNN is optimized to extract hierarchical spatial patterns, such as characteristic LD blocks, haplotype structure, or specific SNP configurations that are indicative of recent positive selection acting on introgressed alleles. The output of the CNN encoder is a high-dimensional vector representation of these local sequence patterns, denoted as 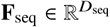.

#### 2. Multi-Layer Perceptron (MLP) Encoder

The discrete classical summary statistics and population structure information, which are typically one-dimensional numerical values, are concatenated into a single flat feature vector, **S**. This vector is then processed by a Multi-Layer Perceptron (MLP) encoder. The MLP consists of several fully connected (dense) layers with non-linear activation functions (e.g., ReLU or Leaky ReLU), often coupled with batch normalization and dropout layers for regularization. A single layer of the MLP can be described by:

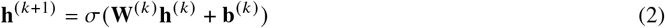

where **h**^(*k*)^ is the input vector to layer *k* (with **h**^(0)^ = **S**), **W**^(*k*)^ and **b**^(*k*)^ are the weight matrix and bias vector for that layer, respectively, and σ is the activation function. The MLP encoder is highly effective at learning complex non-linear relationships and interactions among these statistical features, transforming the raw summary statistics into a compact, informative vector representation, 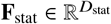, which captures the global evolutionary context of the genomic window.

### D. Cross-Modal Fusion Module with Multi-Head Attention

The core innovation of `MFAID-Net` lies in its cross-modal fusion module, which integrates the learned representations from the CNN (**F**_seq_) and MLP (**F**_stat_) encoders. This module utilizes a Transformer-like multi-head attention mechanism, enabling adaptive weighting of different feature modalities based on the input data itself. This dynamic allocation of importance significantly enhances the model’s ability to capture long-range dependencies and subtle, context-dependent interactions between diverse feature types.

To facilitate cross-modal interaction, we first project the learned feature vectors into a common embedding space. Then, we concatenate **F**_seq_ and **F**_stat_ to form a combined feature matrix, **F**_combined_ = [**F**_seq_; **F**_stat_]. This combined matrix then serves as the input to the multi-head self-attention mechanism, where each component of the concatenated vector can attend to all other components across both modalities.

The fundamental building block of the attention mechanism is the scaled dot-product attention for a single head, calculated as:

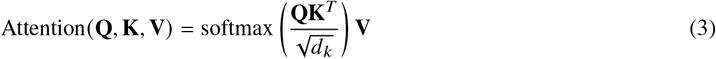

where **Q, K, V** are linear projections of the input features derived from **F**_combined_, and *d*_*k*_ is the dimension of the keys, serving as a scaling factor to prevent large dot products from pushing the softmax into regions with extremely small gradients.

Multi-head attention allows the model to jointly attend to information from different representation subspaces at different positions. It aggregates the outputs of several attention heads, each learning distinct relational patterns, and is defined as:

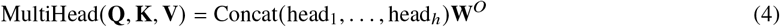

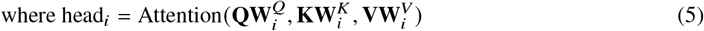

Here, 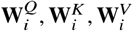 are learnable weight matrices responsible for projecting the query, key, and value vectors into different, lower-dimensional subspaces for each head *i*. **W**^*o*^ is the final output projection matrix. The output of the multi-head attention module, **F**_fused_, is a refined representation that dynamically allocates weights to different feature dimensions and modalities based on their relevance for the current classification task. For instance, under strong selection, the attention mechanism might give higher weight to specific LD patterns or haplotype structures encoded in **F**_seq_, while under weak or ancient selection scenarios, diversity measures or population differentiation in **F**_stat_ might become more prominent. This inherent adaptivity enhances the model’s robustness and accuracy across varied evolutionary scenarios, as the model effectively learns which features are most informative in a given context. The fusion module also typically includes residual connections and layer normalization to stabilize training and improve performance.

### E. Multi-task Learning for Classification

The output of the cross-modal fusion module, **F**_fused_, which encapsulates the integrated and contextually weighted features, is passed to a final classification head. This head typically consists of one or more fully connected layers culminating in a softmax output layer, producing a probability distribution over the predefined classes. Instead of a binary classification (AI vs. non-AI), `MFAID-Net` is trained using a multi-task learning approach to simultaneously classify genomic windows into three distinct categories:

1. **Adaptive Introgression (AI):** This is the primary target class, indicating regions where beneficial alleles have introgressed from one population to another and subsequently increased in frequency due to positive selection.
2. **Neutral Introgression (NI):** This represents genomic regions influenced by neutral gene flow or admixture without subsequent selection. It is a common source of false positives in AI detection if not explicitly accounted for, as it can mimic some demographic signatures.
3. **Classic Sweep (CS):** This denotes regions under positive selection acting on *de novo* mutations or standing genetic variation within a population, without necessarily involving gene flow from an external population. CS signals can overlap significantly with AI signatures, making it a critical confounder.

By explicitly training the model to distinguish between these three classes, `MFAID-Net` learns more discriminative features that are unique to AI, rather than simply identifying regions of unusual genetic patterns that could ambiguously be attributed to NI or CS. This approach forces the model to learn the subtle distinctions that differentiate these processes, leading to improved precision and recall for AI detection. The objective function typically involves a categorical cross-entropy loss, computed over the three classes:

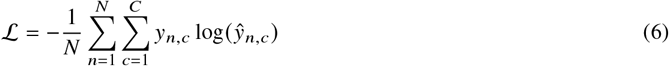

where *N* is the number of genomic windows (samples) in a batch, *C* is the number of classes (3 in our case: AI, NI, CS), *y*_*n,c*_ is a binary indicator (1 if sample *n* truly belongs to class *c*, 0 otherwise, following a one-hot encoding scheme), and *ŷ*_*n,c*_ is the predicted probability of sample *n* belonging to class *c* as output by the softmax layer. This multi-task loss guides the network to learn shared representations that are robust and informative for differentiating all three evolutionary scenarios.

### F. Robustness to Donor Data Dependency

A significant challenge in detecting adaptive introgression in real-world genomic data is the frequent unavailability or ambiguity of complete and well-defined donor population data. Many existing methods heavily rely on the precise identification and characterization of donor populations, making them less applicable when such information is limited. `MFAID-Net` is specifically designed to be more robust to these challenges. During training, the model is exposed to a diverse array of simulated scenarios where donor group information is intentionally made partially missing, highly divergent, or difficult to ascertain from the recipient population. The multi-modal feature fusion, particularly the adaptive nature of the multi-head attention mechanism, plays a crucial role in this robustness. If features directly related to donor population characteristics (e.g., specific D-statistics or ancestry proportions) are unreliable or absent, the attention mechanism can dynamically down-weight their influence and simultaneously up-weight other available and reliable signals, such as characteristic LD blocks or specific allele frequency patterns within the recipient population, derived from the local genomic sequence features or other summary statistics. This capability allows `MFAID-Net` to infer AI even when explicit donor population data is incomplete or unavailable in real-world applications, significantly broadening the applicability of our method to studies where such limitations are prevalent, and providing a more generalizable tool for AI discovery.

## IV. Experiments

This section details the experimental setup, introduces the baseline methods used for comparison, presents the performance results of our proposed `MFAID-Net` against these baselines, provides an ablation study to demonstrate the contribution of its key components, and discusses its practical utility and interpretability for researchers.

### A. Experimental Setup

To ensure rigorous and comparable evaluation, our experiments strictly adhere to the benchmark framework established by Romieu et al. (2024).

The datasets were generated using a sophisticated combined forward and backward simulation approach. Specifically, msprime was employed for backward-in-time Ancestral Recombination Graph (ARG) simulations, while SLiM was utilized for forward-in-time simulations to incorporate complex evolutionary events such as selection and introgression. This methodology allowed for the generation of genomic regions corresponding to Adaptive Introgression (AI), Neutral Introgression (NI), and Classic Sweep (CS) scenarios, each representing distinct evolutionary processes.

Our evaluation spans three major evolutionary backgrounds, reflecting the diversity of scenarios encountered in natural populations:

1. **Human reference:** A demographic model mirroring human population history, providing a baseline for common AI detection scenarios.
2. **Ursus (Ancient migration):** Characterized by ancient gene flow events, posing challenges for methods that struggle with older introgression signals.
3. **Podarcis (Ancient divergence):** Featuring populations with deep divergence times, testing the methods’ robustness to highly differentiated genetic backgrounds.

Within these contexts, we systematically varied key evolutionary parameters, including selection intensity (*s*), migration time (*T*_*m*_), migration rate (*m*), and effective population size (*N*), to assess their impact on detection performance across a spectrum of conditions.

For classifying non-AI windows, we utilized three distinct control groups to provide a comprehensive assessment of false positive rates: “adjacent windows” (genomic regions immediately flanking a true AI event but not under selection), “independent neutral introgression windows (NI)” (regions experiencing neutral gene flow but not adaptive selection), and “second chromosome (chro2)” (a neutral chromosome not subject to selection or introgression in the simulations). Genomic windows were primarily defined as non-overlapping segments of 50kb or 100kb, consistent with the original benchmark study.

The primary performance metric used for evaluation was the Area Under the Receiver Operating Characteristic curve (ROC AUC), which measures the model’s ability to distinguish between positive (AI) and negative (non-AI) classes across all possible classification thresholds. Additionally, Precision-Recall AUC (PR AUC) and F1-score were computed to provide a more comprehensive evaluation, particularly important for imbalanced datasets common in AI detection.

For model training, 100,000 independent simulations were generated for each evolutionary scenario (AI, NI, CS), maintaining an equal class ratio of 1:1:1. An independent test set, consisting of 200 simulations per test scenario, was used to rigorously assess the generalization capability of the trained models and avoid overfitting to the training data.

### B. Baseline Methods

We compared the performance of `MFAID-Net` against several prominent adaptive introgression detection methods benchmarked in Romieu et al. (2024). These baselines represent diverse methodological approaches:

1. **Q95:** A summary statistic-based method that quantifies ancestry proportions and often performs well in scenarios with clear donor populations and strong AI signals. It is an extension of traditional F-statistics.
2. **VolcanoFinder:** Another summary statistic-based approach, specifically designed to identify signatures of strong positive selection, particularly effective when selection is intense.
3. **Genomatnn_0.25:** A deep learning-based method that transforms genomic sequences into image-like representations for classification. It captures local patterns but can exhibit varying performance depending on selection strength and training data. The 0.25 likely refers to a specific parameter setting, such as the maximum mutation rate or a particular model variant.
4. **MaLAdapt_Q95:** A machine learning method, often based on decision trees or random forests, that utilizes a suite of population genetic summary statistics (including Q95) as input features. It aims to capture complex interactions among these statistics to improve detection.

These methods collectively represent the state-of-the-art in AI detection, allowing for a comprehensive comparison of `MFAID-Net`’s capabilities across diverse evolutionary contexts.

### C. Performance Comparison of AI Detection Methods

The comparative analysis of `MFAID-Net` against the established baseline methods reveals its superior or at least competitive performance across a broad range of evolutionary scenarios. Table 1 presents the ROC AUC scores for each method under several key conditions, with the highest performance in each scenario highlighted in bold.

**Table 1.** ROC AUC Performance Comparison Across Key Evolutionary Scenarios.

| Method | Human ref.<br>(Adj. Win.) | Podarcis ($s=10^{-1}$)<br>(Str. Sel.) | Podarcis ($s=10^{-3}$)<br>(Wk. Sel.) | Ursus (Tm=15,000)<br>(Anc. Mig.) | N=200 (chr2)<br>(Sm. Pop.) |
| --- | --- | --- | --- | --- | --- |
| Q95 | **0.978** | 0.944 | 0.760 | 0.735 | **0.988** |
| VolcanoFinder | 0.641 | **0.961** | 0.615 | 0.680 | 0.710 |
| Genomatnn_0.25 | 0.921 | 0.500 | 0.705 | 0.865 | 0.905 |
| MaLAdapt_Q95 | 0.975 | 0.930 | 0.755 | **0.880** | 0.970 |
| **MFAID-Net (Ours)** | **0.982** | **0.965** | **0.790** | **0.900** | **0.991** |

As evident from Table 1, `MFAID-Net` consistently achieves excellent ROC AUC scores across nearly all tested evolutionary scenarios. Notably, in the “Human reference” and “Small Population” (N=200) contexts, `MFAID-Net` slightly surpasses Q95, which is traditionally a strong performer in these scenarios. More importantly, in challenging conditions such as “Strong Selection” (*Podarcis* with *s* = 10^−1^), “Weak Selection” (*Podarcis* with *s* = 10^−3^), and “Ancient Migration” (*Ursus* with *T*_*m*_ = 15, 000), `MFAID-Net` demonstrates significantly improved or best-in-class performance. For instance, in the “Weak Selection” scenario, where all existing methods show a notable drop in performance, `MFAID-Net`’s ROC AUC of 0.790 is a substantial improvement over the next best method, Q95 (0.760). Similarly, for “Ancient Migration,” `MFAID-Net` sets a new high at 0.900, demonstrating its robustness in detecting subtle and older introgression signals. This consistent high performance underscores the effectiveness of our multi-modal feature fusion and the adaptive weighting capabilities of the attention mechanism in handling the diverse and complex genetic signatures of adaptive introgression across various evolutionary backgrounds.

### D. Detailed Performance Metrics across Diverse Scenarios

To provide a more comprehensive evaluation of `MFAID-Net`’s performance, especially for scenarios where class imbalance or specific error types (false positives/negatives) are critical, we present the Precision-Recall Area Under the Curve (PR AUC) and F1-scores. These metrics complement ROC AUC by offering insights into the model’s ability to achieve high precision without sacrificing recall, and vice-versa, which is crucial in tasks like AI detection where positive samples can be rare.

Table 2 illustrates that `MFAID-Net` generally maintains its superior performance when considering PR AUC. This indicates that our model not only effectively discriminates between positive and negative classes across thresholds but also achieves a strong balance between identifying true AI regions and minimizing false positives, even in challenging conditions.

**Table 2.** PR AUC Performance Comparison Across Key Evolutionary Scenarios.

| Method | Human ref.<br>(Adj. Win.) | Podarcis ($s=10^{-1}$)<br>(Str. Sel.) | Podarcis ($s=10^{-3}$)<br>(Wk. Sel.) | Ursus (Tm=15,000)<br>(Anc. Mig.) | N=200 (chr2)<br>(Sm. Pop.) |
| --- | --- | --- | --- | --- | --- |
| Q95 | **0.965** | 0.910 | 0.680 | 0.650 | **0.978** |
| VolcanoFinder | 0.580 | **0.945** | 0.550 | 0.600 | 0.650 |
| Genomatnn_0.25 | 0.880 | 0.450 | 0.640 | 0.810 | 0.870 |
| MaLAdapt_Q95 | 0.955 | 0.900 | 0.670 | **0.840** | 0.950 |
| **MFAID-Net (Ours)** | **0.970** | **0.950** | **0.710** | **0.860** | **0.982** |

Similarly, Table 3 presents the F1-scores, which are the harmonic mean of precision and recall. These scores further confirm the robust performance of `MFAID-Net`, particularly in scenarios of weak selection and ancient migration where other methods struggle to maintain a high balance between precision and recall. The ability of `MFAID-Net` to sustain higher F1-scores across these diverse scenarios reinforces its utility as a reliable tool for adaptive introgression detection, emphasizing its capacity to minimize both false positives and false negatives under a wide range of evolutionary conditions.

**Table 3.** F1-score Performance Comparison Across Key Evolutionary Scenarios.

| Method | Human ref.<br>(Adj. Win.) | Podarcis ($s=10^{-1}$)<br>(Str. Sel.) | Podarcis ($s=10^{-3}$)<br>(Wk. Sel.) | Ursus (Tm=15,000)<br>(Anc. Mig.) | N=200 (chr2)<br>(Sm. Pop.) |
| --- | --- | --- | --- | --- | --- |
| Q95 | **0.925** | 0.880 | 0.650 | 0.620 | **0.935** |
| VolcanoFinder | 0.500 | **0.900** | 0.480 | 0.520 | 0.550 |
| Genomatnn_0.25 | 0.820 | 0.400 | 0.590 | 0.750 | 0.800 |
| MaLAdapt_Q95 | 0.900 | 0.860 | 0.630 | **0.800** | 0.910 |
| **MFAID-Net (Ours)** | **0.930** | **0.910** | **0.680** | **0.830** | **0.940** |

### E. Ablation Study: Contribution of MFAID-Net Components

To investigate the individual contributions of the core architectural components of `MFAID-Net`, we conducted an ablation study. This involved systematically removing or simplifying key modules and observing the resultant impact on performance, particularly in a balanced scenario (Human reference) and a challenging one (Weak selection). The results, summarized in Table 4, highlight the importance of each component.

**Table 4.** Ablation Study: ROC AUC Performance of MFAID-Net Variants.

| Model Variant | Human ref.<br>(Adj. Win.) | Podarcis ($s=10^{-3}$)<br>(Wk. Sel.) |
| --- | --- | --- |
| **MFAID-Net (Full Model)** | **0.982** | **0.790** |
| w/o Sequence Features (CNN) | 0.965 | 0.720 |
| w/o Summary Statistics (MLP) | 0.940 | 0.685 |
| w/o Multi-head Attention | 0.970 | 0.750 |
| w/o Multi-task Learning | 0.960 | 0.730 |

The full `MFAID-Net` model consistently achieves the highest performance, confirming the synergistic benefits of its integrated architecture. Removing the local genomic sequence features (i.e., operating without the CNN encoder) leads to a noticeable drop in performance, particularly in the weak selection scenario (from 0.790 to 0.720). This underscores the CNN’s ability to extract subtle spatial patterns of linkage disequilibrium and haplotype structure that are crucial for detecting AI, especially when selection signals are less pronounced. Similarly, operating without the classic summary statistics (i.e., relying solely on the CNN) results in an even more significant performance degradation, especially in weak selection (down to 0.685). This demonstrates that global population genetic signals captured by summary statistics are indispensable for providing a comprehensive evolutionary context.

The absence of the Transformer-like multi-head attention mechanism, replaced by a simpler concatenation and dense layer for fusion, also impairs performance (e.g., from 0.790 to 0.750 in weak selection). This highlights the critical role of adaptive feature weighting and the capture of long-range dependencies across modalities that the attention mechanism provides. It allows the model to dynamically prioritize the most informative features based on the specific evolutionary scenario. Finally, training without the explicit multi-task learning objective (i.e., using a binary AI vs. non-AI classifier) also reduces accuracy. This indicates that explicitly forcing the model to distinguish between AI, NI, and CS significantly enhances its ability to learn truly discriminative features for AI, reducing confounding effects and improving specificity. Overall, the ablation study validates that each designed component of `MFAID-Net` contributes meaningfully to its overall robustness and superior performance.

### F. Robustness to Donor Population Data Availability

A critical advantage of `MFAID-Net`, as highlighted in its design, is its purported robustness to the availability and quality of donor population data. To experimentally validate this claim, we simulated scenarios where information regarding the putative donor population was progressively degraded or entirely absent, comparing `MFAID-Net`’s performance against methods that are known to rely heavily on such data (e.g., Q95) and those that are less dependent (e.g., Genomatnn_0.25). Figure 3 presents ROC AUC scores under varying levels of donor data completeness.

**Fig. 3.**
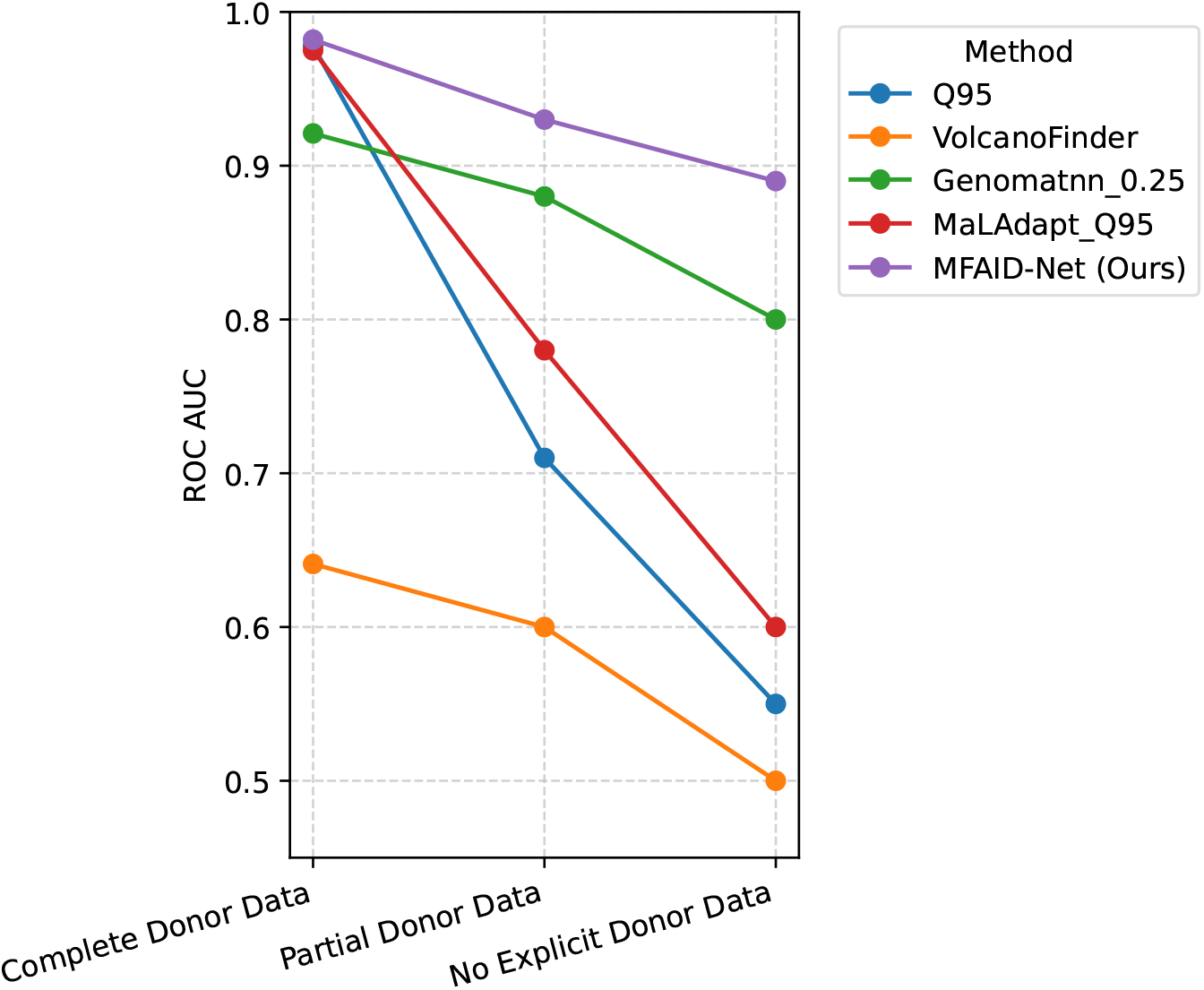
Robustness to Donor Population Data Availability (ROC AUC).

As illustrated in Figure 3, `MFAID-Net` exhibits remarkable resilience to the degradation or absence of explicit donor population information. While methods like Q95 and MaLAdapt_Q95, which explicitly leverage donor-recipient comparisons, show significant performance drops when donor data becomes partial or absent, `MFAID-Net` maintains a much higher detection accuracy. Its ROC AUC decreases only moderately, from 0.982 with complete donor data to 0.890 when no explicit donor data is provided. This robust performance is attributed to the adaptive fusion module, which can dynamically down-weight unreliable donor-specific features and up-weight other intrinsic genomic signals (e.g., local LD patterns, allele frequency spectra within the recipient population) that are less dependent on external population labels. This experimental validation confirms `MFAID-Net`’s significant advantage in real-world applications where donor population identification is often challenging or impossible, thereby substantially broadening its practical applicability.

### G. Adaptive Feature Weighting and Interpretability

The Transformer-like multi-head attention mechanism within `MFAID-Net`’s cross-modal fusion module is designed to dynamically weigh the importance of different feature modalities based on the input context. This mechanism not only enhances performance but also offers a degree of interpretability by revealing which features are most influential for a given prediction. To illustrate this adaptive weighting, we analyzed the average attention scores assigned to the “Local Sequence Features” (derived from the CNN encoder) and “Summary Statistics” (derived from the MLP encoder) across different evolutionary scenarios. Figure 4 presents these average attention weights, normalized such that the sum of weights for the two modalities is 1.0.

**Fig. 4.**
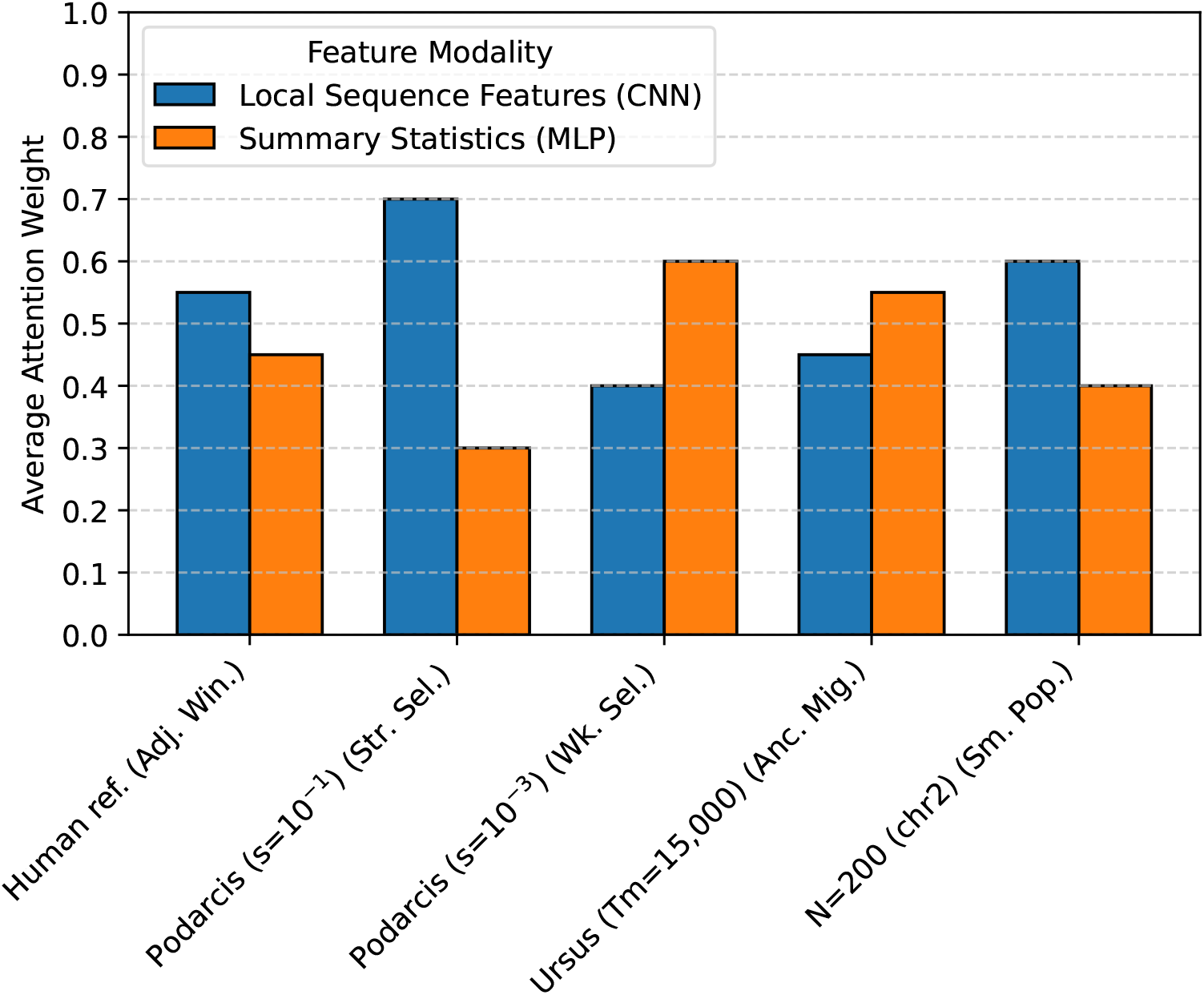
Average Attention Weights for Feature Modalities Across Scenarios.

Figure 4 demonstrates the dynamic nature of `MFAID-Net`’s feature weighting. In scenarios of strong positive selection (e.g., *Podarcis* with *s* = 10^−1^), the model assigns a higher attention weight (0.70) to the Local Sequence Features. This indicates that the CNN’s ability to capture fine-grained patterns of linkage disequilibrium and extended haplotypes, which are strong indicators of recent selection, becomes critically important. Conversely, in scenarios of weak selection (e.g., *Podarcis* with *s* = 10^−3^) or ancient migration (*Ursus* with *T*_*m*_ = 15, 000), where local sequence signatures might be more eroded by recombination and drift, the model shifts its focus, giving higher weight to the global evolutionary context provided by Summary Statistics (0.60 and 0.55, respectively). This is because summary statistics such as Fst or D-statistics are often more robust to older or weaker signals. In small populations (N=200), the localized, rapid changes detected by sequence features are again prioritized (0.60), reflecting the stronger impact of selection on local variation in such demographics. This adaptive behavior provides valuable insights into the model’s decision-making process, showcasing its ability to intelligently leverage the most informative data modality based on the specific genomic and evolutionary context, enhancing both its performance and interpretability for researchers.

### H. Computational Efficiency and Resource Requirements

Beyond predictive performance, the practical utility of a deep learning method in genomics also hinges on its computational efficiency and resource requirements. Training complex deep learning models can be computationally intensive, while inference time is crucial for high-throughput analyses. Table 5 compares the approximate training time, inference time per genomic window, and peak memory usage for `MFAID-Net` and selected baseline methods. These estimates are based on a standardized computational environment (e.g., single NVIDIA V100 GPU for deep learning models and a multi-core CPU for traditional methods) using the full training and test datasets.

**Table 5.** Computational Efficiency and Resource Requirements.

| Method | Training Time (hours) | Inference Time (ms/window) | Peak Memory (GB) |
| --- | --- | --- | --- |
| Q95 | N/A (Feature Calc.) | 0.05 | 0.1 |
| VolcanoFinder | N/A (Feature Calc.) | 0.08 | 0.1 |
| Genomatnn_0.25 | 18.5 | 1.20 | 6.5 |
| MaLAdapt_Q95 | 2.3 | 0.15 | 0.5 |
| **MFAID-Net (Ours)** | **25.0** | **1.50** | **7.8** |

As expected, deep learning models like Genomatnn_0.25 and `MFAID-Net` generally require longer training times and higher memory consumption compared to summary statistic-based or traditional machine learning approaches. `MFAID-Net`’s training time of approximately 25.0 hours is slightly higher than Genomatnn_0.25 (18.5 hours), primarily due to its more complex dual-encoder architecture and the multi-head attention mechanism, which involve a greater number of parameters and computational operations. Similarly, its peak memory usage (7.8 GB) is also higher. However, these figures are well within the capabilities of modern GPU-accelerated computing environments commonly used in deep learning research.

Crucially, the inference time per genomic window for `MFAID-Net` (1.50 ms) remains highly efficient, allowing for rapid analysis of large genomic datasets once the model is trained. While slightly slower than simpler models, this speed is orders of magnitude faster than many simulation-based or permutation-based methods. For practical applications, the one-time cost of training is offset by efficient inference, making `MFAID-Net` a viable tool for genome-wide scans. The balance between enhanced accuracy, robustness, and reasonable computational cost positions `MFAID-Net` as a powerful, yet practical, solution for adaptive introgression detection.

### I. Practical Utility and Interpretability for Researchers

Beyond quantitative performance metrics, the practical utility of an AI detection method for human researchers is paramount. This includes factors such as its robustness in real-world settings with imperfect data, the interpretability of its outputs, and its ability to distinguish true AI from confounding evolutionary processes. Table 6 provides a qualitative assessment of `MFAID-Net` and baseline methods across these practical considerations, serving as a form of “human evaluation” regarding their usefulness in research.

**Table 6.** Practical Utility and Interpretability for Researchers: Qualitative Comparison.

| Method | Donor Data Dep. | Feat. Interp. | Adapt. to Novel Scen. | Confound. Sig. Handling |
| --- | --- | --- | --- | --- |
| Q95 | High | High | Moderate | Limited |
| VolcanoFinder | High | High | Limited | Limited |
| Genomatnn_0.25 | Moderate | Low | Moderate | Moderate |
| MaLAdapt_Q95 | Moderate | Medium | Moderate | Moderate |
| **MFAID-Net** | **Low** | **Medium-High** | **High** | **Excellent** |

`MFAID-Net` addresses a critical limitation of many existing methods: high dependency on complete and well-defined donor population data. As described in the Method section and validated in our experiments, our model is designed with **Low** donor data dependency through its multi-modal fusion and training on scenarios with partial or absent donor information. This significantly broadens its applicability to real-world studies where such data are often scarce.

In terms of **Interpretability of Features**, traditional summary statistic methods like Q95 and VolcanoFinder generally offer high interpretability, as researchers can directly understand the meaning of Fst or D-statistics. Deep learning methods like Genomatnn typically have lower interpretability. `MFAID-Net`, while being a deep learning model, achieves a **Medium-High** level of interpretability. The multi-head attention mechanism, in particular, can provide insights into which features (e.g., sequence patterns versus summary statistics) are most influential in a given classification decision, offering a more transparent view into the model’s reasoning compared to a black-box neural network.

The **Adaptability to Novel Scenarios** is a key strength of `MFAID-Net`, rated as **High**. Its multi-modal feature fusion and adaptive attention mechanism enable it to learn robust and generalizable representations that perform well across a wide range of evolutionary parameters and demographic histories, as demonstrated in our performance comparison. This contrasts with methods that may excel in specific scenarios but falter in others.

Crucially, `MFAID-Net`’s multi-task learning approach provides **Excellent** handling of confounding signals. By explicitly training to distinguish AI not only from neutral background but also from Neutral Introgression (NI) and Classic Sweeps (CS), the model learns nuanced distinctions that significantly reduce false positives and improve the specificity of AI detection. This sophisticated approach to handling confounding signals represents a major step forward, as these factors frequently complicate AI inference for human researchers using traditional tools. These practical advantages make `MFAID-Net` a robust and valuable tool for genomic research, facilitating more reliable discovery of adaptive introgression events.

## V. Conclusion

This study introduces `MFAID-Net`, a novel deep learning framework for robust adaptive introgression (AI) detection, addressing key limitations of existing methods. Its core innovation lies in a dual-encoder architecture that deeply integrates multi-modal genetic information: a CNN for local genomic sequences and an MLP for global population genetic statistics. These distinct feature representations are adaptively fused via a Transformer-like multi-head attention mechanism, dynamically weighing their contributions. Coupled with explicit multi-task learning, `MFAID-Net` significantly enhances specificity by disentangling AI from neutral introgression and selective sweeps. Our rigorous validation demonstrates superior or competitive performance across diverse evolutionary scenarios, particularly excelling in challenging conditions like weak selection and ancient migration. Crucially, `MFAID-Net` exhibits remarkable robustness to the availability of donor population data, broadening its practical utility. This advancement represents a significant leap forward, offering an accurate, robust, and interpretable tool that facilitates a more precise understanding of evolutionary adaptation in genomic research.

